# Phylo-comparative analyses reveal the dual role of drift and selection in reproductive character displacement

**DOI:** 10.1101/610857

**Authors:** İsmail K. Sağlam, Michael R. Miller, Sean O’Rourke, Selim S. Çağlar

**Affiliations:** Koç University, Department of Molecular Biology and Genetics, Istanbul, Turkey; University of California Davis, Department of Animal Science, Davis, CA, USA; Hacettepe University, Department of Biology, Ankara, Turkey

**Keywords:** Quaternary climate shifts, phenotypic trajectories, evolutionary rates, selective regimes, *Phonochorion*, Caucasus

## Abstract

When incipient species meet in secondary contact, natural selection can rapidly reduce costly reproductive interactions by directly targeting reproductive traits. This process, called reproductive character displacement (RCD), leaves a characteristic pattern of geographic variation where divergence of traits between species is greater in sympatry than allopatry. However, because other forces can also cause similar patterns, care must be given in separating pattern from process. Here we show how the phylo-comparative method together with genomic data can be used to evaluate evolutionary processes at the population level in closely related species. Using this framework, we test the role of RCD in speciation of two cricket species endemic to Anatolian mountains by quantifying patterns of character displacement, rates of evolution and adaptive divergence. Our results show differing patterns of character displacement between species for reproductive vs. non-reproductive characters and strong patterns of asymmetric divergence. We demonstrate diversification results from rapid divergence of reproductive traits towards multiple optima under the dual influence of strong drift and selection. These results present the first solid evidence for RCD in Anatolian mountains, quantify the amount of drift and selection necessary for RCD to lead to speciation, and demonstrate the utility of phylo-comparative methods for quantifying evolutionary parameters at the population level.

## 1. Introduction

Reproductive character displacement (RCD), is the process where divergence in reproductive traits is driven directly by selection in order to reduce costly reproductive interactions between incipient species (Pfennig and Pfennig, 2009). When hybridization is involved, the process can also be referred to as reinforcement (Ortiz-Barrientos *et al*., 2009; Servedio, 2004). RCD leaves a characteristic pattern of geographic variation where phenotypic divergence of reproductive traits is greater between species in sympatry than allopatry (Jang and Gerhardt, 2007; Jang et al., 2009; Kameda et al., 2009; Lemmon, 2009). However, concluding RCD from patterns of phenotypic variation is misleading because other forces can also lead to similar results (Templeton, 1981; Day, 2000; Vallin et al., 2011). For example, greater divergence of reproductive traits in sympatry can often result as a by-product of ecological character displacement (ECD, divergence between species due to resource competition) (Podos and Nowicki, 2004; Huber and Podos, 2006). Therefore, care must be given in separating pattern from process.

RCD is especially important in zones of secondary contact between species that were previously isolated but did not have sufficient time to reach full prezygotic isolation (Zamudio and Savage, 2003; Noonan and Gaucher, 2006; Dubey et al., 2008). In the absence of strong prezygotic barriers, the production of maladaptive hybrids can result in strong selection for rapid divergence of reproductive traits (Servedio, 2004a, 2004b; Stewart et al., 2016). For example, climatic oscillations during the Quaternary and subsequent range shifts of organisms could have created numerous opportunities for secondary contact between partially diverged lineages (Stewart et al., 2013). With frequent distributional shifts in response to glacial cycles, there presumably would have been multiple opportunities for population isolation and divergence, but also for gene flow when partially diverged gene pools came into contact during range expansions (Jansson and Dynesius, 2002; Knowles and Carstens, 2007; Galbreath et al., 2010). In theory, this would present a suitable medium for a process like RCD (reinforcement or otherwise) to act (Servedio, 2004a; Stewart, 2013). Phylogeographic studies have documented numerous secondary contact zones resulting from the effects of glacial cycles (Zamudio and Savage, 2003; Noonan and Gaucher, 2006; Alberto et al., 2008; Ruiz et al., 2012). However, information on the mechanisms producing/maintaining diversification in these systems or how recurring fragmentation and admixture might have influenced reproductive isolation is notably missing (Stewart, 2013).

Species heavily influenced by glacial cycles are also characterized by low overall diversity, high amount of genetic fragmentation and strong bottlenecks (Knowles, 2001a; b; Knowles and Richards, 2005; Kaya and Çıplak 2016). All these factors can greatly obscure or erase past signals of evolutionary processes, especially when coupled with strong diversifying selection as presumed by RCD. Indeed, RCD has mostly been observed from stable hybrid zones, where the action of past processes leading to speciation can be observed in a contemporary setting, or in organisms which can actively hybridize either in the wild or in the laboratory (Hoskin et al., 2005; Nosil et al., 2007; Latour et al., 2014; Stewart et al., 2013). In the absence of stable hybrid zones, quantifying the effect of RCD on the speciation process is extremely difficult, especially when direct experimental manipulation like laboratory crosses are not possible.

When contemporary populations do not present a suitable medium for measuring the effect of past selective pressures, the phylo-comparative method can be a powerful substitute (Freckleton et al., 2011). By using natural variation within and between species along a phylogeny, the comparative method has emerged as an effective tool for testing and quantifying past ecological and evolutionary processes (Felsenstein, 1985, 1988; Garland et al., 1992; Hey and Nielsen, 2007; Smith, 2008; Freckleton and Jetz, 2009). This approach can now infer the presence and effect of different selective optima (Butler and King, 2004), detect and compare rates of trait evolution (Adams, 2013), detect correlated evolution among traits (Pagel and Meade, 2006), and trace shifts in trait correlation through evolutionary time (Eastman et al., 2011; Revell et al., 2011).

In this study, we use genome wide data together with the phylo-comparative method to show the utility of these methods for quantifying evolutionary processes such as RCD in natural populations. We use two species of montane bush crickets - *Phonochorion uvarovi* Uvarov, 1916 and *Phonochorion artvinensis* Bei-Bienko, 1954 (Orthoptera: Phaneropterinae) - endemic to the Lesser Caucasus mountains of Northeastern Turkey (Fig. 1) (Sevgili et al., 2010; Sağlam et al., 2014). Anatolian mountains are well known biodiversity hotspots repeatedly influenced by glacial cycles (Çıplak, 2004; Bilgin et al., 2006; Şekercioğlu et al., 2011; Gür, 2013; Özüdoğru et al., 2015; Perktaş et al., 2015). Moreover, several reviews (Bilgin, 2011; Çıplak, 2008) propose speciation via hybrid unfitness as a dominant mechanism in producing diversity along Anatolian mountains, either within potential hybrid tension zones (Bilgin, 2011), or spanning from repeated admixture of populations during elevation shifts (Çıplak, 2008). Recent studies conducted on high altitude endemics of Anatolia reveal numerous cases where divergence patterns are highly suggestive of RCD (Sağlam et al., 2014; Çıplak et al., 2015; Kaya et al., 2015), however explicit tests are yet to be performed.

**Fig. 1.**
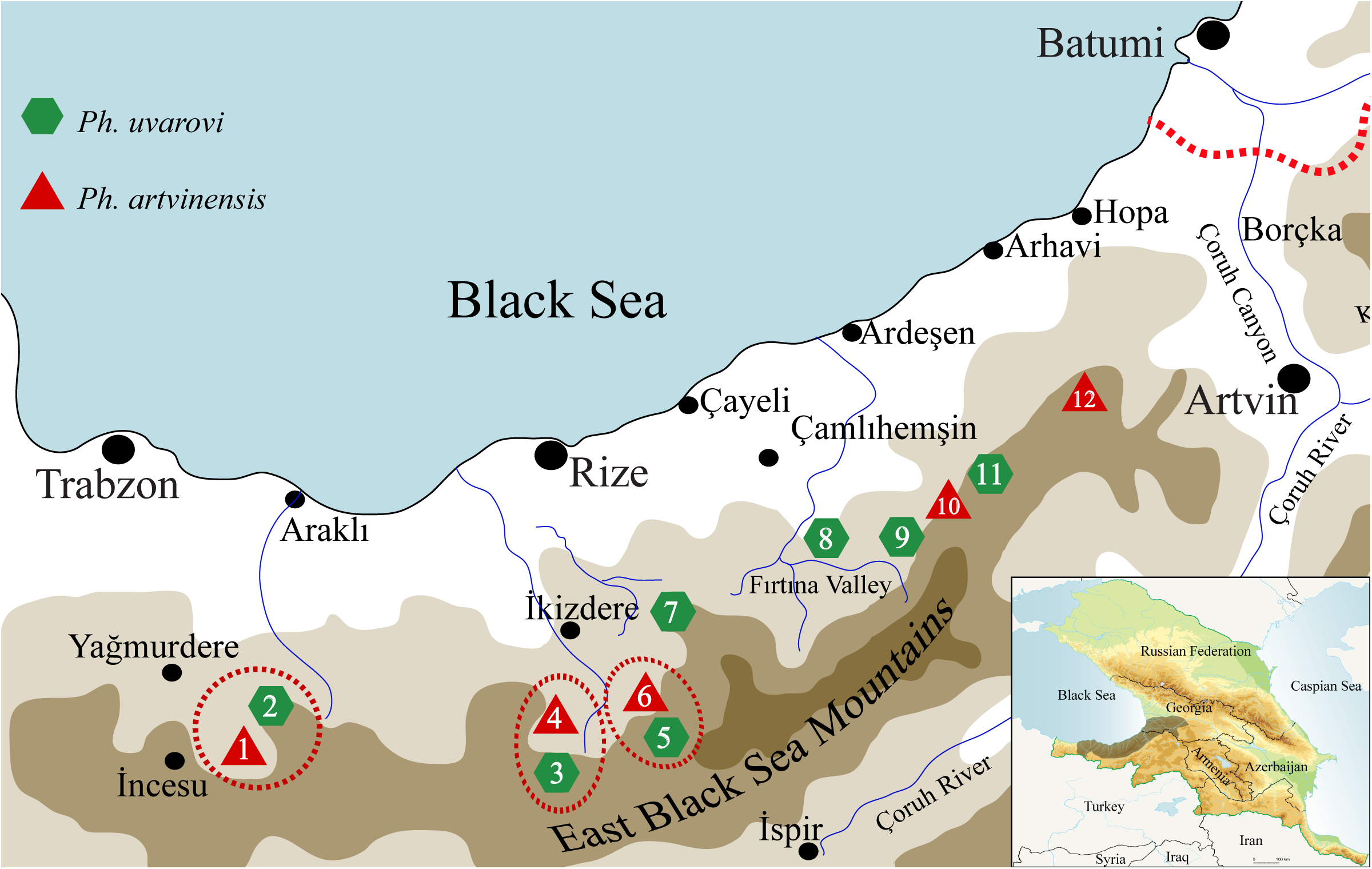
Distribution of Phonochorion species within the study area. Populations used within the have been numbered and details can be found in Table S1 by following the corresponding numbers. Dashed circles correspond to sympatric populations whereas all other populations are allopatric.

To quantify the role of RCD in divergence and speciation of *Ph. artvinensis* and *Ph. uvarovi* we reconstructed the evolutionary history of allopatric and sympatric populations of the two species using two independent phylogenetic sets. Each set was made up of 50 loci chosen from a genomic library constructed using RAD sequencing, and ITS and mtDNA sequences. From the estimated phylogenies we then inferred the tempo and mode of trait divergence for two male traits which are hypothesized to give different responses to selection for reproductive isolation: 1) the subgenital plate, a reproductive trait which can rapidly respond to selection for reproductive isolation as it plays an active role in spermatophore transfer and hence species specificity (Heller et al., 2004; Helversen and Helversen, 2004) and 2) the pronotum, a morphological character not directly under selection for reproductive isolation but a trait known to respond to a host of ecological factors (Bidau and Martí, 2007, 2008; Whitman, 2008; Çağlar et al., 2014).

Using the phylo-comparative framework we set out to quantify and test three specific predictions of the role of RCD in diversification/speciation: 1) Greater divergence in sympatry vs. allopatry should only be observed in the reproductive trait; 2) Reproductive characters should show faster rates of divergence and; 3) Divergence of reproductive traits should fit selection models with two selective optima representing differential divergence in sympatry vs. allopatry. By testing these predictions, we reveal that current patterns of RCD between allopatric and sympatric populations of bush cricket species resulted from rapid divergence of reproductive traits towards multiple optima under both strong drift and selection. These results demonstrate the usefulness of the phylo-comparative method for quantifying evolutionary hypotheses at the population level and provide insight regarding the amount of drift and selection necessary for RCD to lead to speciation in a relatively short time.

## 2. Materials and methods

### 2.1. Sampling

Analyses were based on 12 populations chosen from previous field surveys (Sağlam et al., 2014). Distribution of populations and species affiliation are given in Fig. 1. All populations were classified as either sympatric or allopatric (Supplementary material, Table S1). Populations situated in different valley systems and trapped in sky islands (7, 8, 9, 10, 11 and 12, Fig. 1) were classified as allopatric while populations found in the same valley system were classified as sympatric (1, 2, 3, 4, 5 and 6, Fig. 1). Within the zone of sympatry species contain partially isolated populations as *Ph. uvarovi* individuals are mostly distributed at higher altitudes (1800 – 2000 meters) while *Ph. artvinensis* individuals are distributed at lower altitudes (1600 – 1800 meters). However around 1700 – 1800 meters, the two species can be found in true sympatry (Fig. 1). In total we analysed 3 sympatric populations of *Ph. uvarovi* and *Ph. artvinensis*, 4 allopatric populations of *Ph. uvarovi* and 2 allopatric populations of *Ph. artvinensis*. Population information, number of samples used in analyses, and mean values of measured traits are given in Table S1.

### 2.2. Analysis of shape

To quantify morphological divergence in the subgenital plate (SGP) and pronotum (PRNT) of males, shape variation was summarized using elliptic fourier descriptors (Ferson et al., 1985; Rohlf, 1990). Here, *x* and *y* coordinates tracing the outline of traits (Fig. S1) are fit separately as functions of arc length by fourier transformation and later summarized into principal components (PCs). PCs can be considered as reorganized, uncorrelated morphological traits representing different aspects of total shape variation (Fig. S2). All computations were done in SHAPE package v1.2 (Iwata and Ukai, 2006).

### 2.3. Sequence generation and phylogenetic data sets

Sequence data used in this study came from: 1) a partial *de-novo* genomic reference of 45,039 RAD-contigs (length > 300 bp) generated by a new RAD protocol (Ali et al., 2016); 2) complete sequences of the ITS1 and ITS2 regions; and 3) from a 1002 bp concatenated mtDNA data set comprised of the ND4 (582 bp), Cytb (312 bp) and ND1 (108 bp) genes. For details concerning DNA extraction, library preparation, amplification and sequencing see supplementary material. Phylogenetic reconstructions were carried out using two independent genomic subsets, each made up of 50 RAD-loci, randomly chosen from our set of reference RAD-contigs. To each 50 RAD-loci set we further added all available mtDNA and ITS sequences of each population to produce two phylogenetic data sets, each comprised of 52 loci. Working with two independent sets of random genomic loci enabled us to test the robustness of our phylogenetic estimates. Independence of phylogenetic sets were robust to the inclusion of shared mtDNA and ITS regions as discussed fully in supplementary material.

### 2.4. Phylogenetic analyses

Phylogenetic relationships between populations were estimated separately for the two loci sets using the multi-species coalescent procedure (*BEAST) (Heled and Drummond, 2010) as implemented in the program BEAST v.2.2 (Bouckaert et al., 2014). The multi-species coalescent uses the variance in gene genealogies nested within species trees to accurately describe species transmissions (i.e. species divergence history) (Drummond and Boukaert 2015). Here we used the same procedure to describe the phylogenetic relationship between sampled populations. Hence our goal was to estimate population divergence history as correctly as possible from gene genealogies nested within each population.

All analyses were set up using a strict molecular clock, a linear population size model for the multi-species coalescent and a Yule model of divergence for populations (additional information given in supplementary material). Trees were calibrated by setting the mtDNA substitution rate to 0.0133 ± 0.0013 (subs/s/my/l) corresponding to a global mtDNA rate of 2.6% divergence per myr. The effect of different substitution rates and relaxed clock models on divergence times were discussed in another paper (Sağlam et al., 2014) and readers are kindly referred to that paper and supplementary material for further information.

For each set of loci we conducted two independent MCMC runs with chain lengths of 100 million, sampling every 10,000 generations. 10,000 trees were sampled from each run, 25% of which were discarded as burn-in, and joined using LogCombiner v1.7.1 to give us a final sample size of 15,000 trees for each subset. Consensus trees with divergence times were estimated for each subset using TreeAnnotator v.2.2, and visualized using FigTree v.4 (http://tree.bio.ed.ac.uk/software/figtree) and DensiTree 2 (Bouckaert and Heled, 2014).

### 2.5. Character Displacement

To determine patterns of character displacement between species we performed a two-way MANCOVA. Significant PC axes summarizing shape variation were used as dependent variables, “species” and “type” (allopatric vs. sympatric) were entered as independent factors and “size” and “altitude” were entered as covariates. Here a significant “species*type” interaction is indicative of character displacement. To determine whether patterns persisted when controlled for shared ancestry we conducted a phylogenetic MANCOVA (pMANCOVA) using mean PC values, the consensus population tree and the mvgls and manova.gls functions available in the developmental version of mvMORPH (Clavel et al., 2015, 2019). The same linear model used in the traditional two-way MANCOVA was fitted using generalized least squares under Pagel’s lambda transformation and the Mahalanobis method which is an approximation of the leave one out cross-validation of the log-likelihood. MANOVA was performed on the fitted model using Wilk’s lambda statistics and significance was computed using 999 permutations.

We also calculated the degree of divergence of species in allopatry (*D*_*allo*_) and sympatry (*D*_*symp*_) using “average squared mahalanobis distance” between populations. Mahalanobis distance between populations (i.e. distance between population centers in principal component space accounting for different variances) was calculated using the “maha” function in the R package GenAlgo (https://rdrr.io/cran/GenAlgo/man/maha.html). Statistical significance of *D*_*symp*_ *– D*_*allo*_ was determined through a randomization test with 999 iterations where the probability of finding a more extreme value than the observed was the significance level (*P*_*rand*_).

### 2.6. Phenotypic trajectories

To investigate whether character displacement resulted from the magnitude and/or direction of phenotypic change we used phenotypic trajectory analysis (Adams and Collyer, 2009) to calculate phenotypic change vectors (PCVs) between species at two levels: allopatry and sympatry. Statistical significance between the angle (direction) and length (magnitude) of PCVs were assessed by residual randomization (Collyer and Adams, 2007). The difference in the magnitude of phenotypic change reflects whether one species exhibits greater differentiation relative to the other (asymmetric divergence), while the direction of change can be used to assess patterns of convergence, divergence, or parallelism of trait evolution (Adams and Collyer, 2007, 2009).

### 2.7. Divergence rate of traits

To investigate rates of trait evolution we compared the likelihood of a model where each trait evolves at a distinct rate along a given phylogeny with a model where all traits are constrained to evolve at a common rate. Rates of trait evolution were calculated using a Brownian model of evolution and time calibrated phylogenetic trees using the procedure in Adams (2013). To take phylogenetic uncertainty and the effect of different loci sets into account, analyses were conducted separately for both phylogenetic data sets using all of the 15,000 post burn-in trees. For each trait, mean population values of PC1 scores were used as input data along with intra-population variation (i.e. measurement error) (Table S1). The fit of the two different models (i.e. unconstrained vs. constrained rate) was compared using a likelihood ratio test. All analyses were performed in R version 3.0.2 (Team, 2013) using the APE library (Paradis et al., 2004, 2014) and the R code given in Adams (2013).

### 2.8. Comparing neutral and adaptive models of trait evolution

To test if the degree of trait divergence between species in sympatry vs. allopatry was adaptive in nature, we compared neutral (Brownian motion, BM) and adaptive (Ornstein-Uhlenbeck, OU) models of trait divergence as given in Butler and King (Butler and King, 2004) using the R package OUCH (King et al., 2009). Three alternative models were constructed by assigning hypothetical selective regimes to the internal branches of the phylogeny (Fig. S3): 1) the drift only model (DM) where none of the branches were assigned a selective optima; 2) the single adaptive peak model (SAP) where a single (global) optima was assigned to all branches; and 3) the double adaptive peak model (DAP) where two distinct optima designated as “average” and “high” were assigned to allopatric and sympatric populations respectively. Mahalanobis distances were used to calculate the average distance an allopatric population of one species had from allopatric populations of the other species and vice versa for sympatric populations. The resulting mean values for each population (Table S2) were used as continuous variables in all analyses. Analyses were conducted separately for both phylogenetic data sets using all of the 15,000 post burn-in trees. The fit of different models to the data was assessed using likelihood scores and for each model we also calculated optimum trait values (θ_*i*_) and coefficients of drift (σ) and selection (α).

## 3. Results

### 3.1. Shape variation

The total number of principal components describing a significant portion of shape variation were 4 and 9 for the subgenital plate and pronotum respectively (Table S3). In total, the significant PC axes explained over 95% of the variation in both traits with the first PC axes explaining approximately 80 and 46% of the total variation (Table S3 and Fig. S2). We conclude that fourier analysis was able to effectively capture shape variation in both traits.

### 3.2. Robust phylogenetic relationships under different loci sets

Branch lengths and topology of trees reconstructed from each of the two loci sets were consistent with each other (Fig. 2A), so we summarized the entire tree space into a single consensus tree (Fig. 2B). Separation between species was clear as populations of both species clustered into well-supported monophyletic clades. For *Ph. artvinensis*, phylogenetic relationships between populations were also well resolved as posterior probabilities of nodes were high. However, relationships between populations of *Ph. uvarovi* were much less clear (Fig. 2). Divergence times estimated from both loci sets were extremely close and indicated that the two species split fairly recently between 60 – 190 ma (Table 1). The high similarity in topology and branch lengths obtained from the two phylogenetic sets strongly suggests we reliably captured the evolutionary history of populations. We conclude that divergence between species and populations evolved fairly recently.

**Table 1.** Divergence time of species and timing of population splits inferred using the multi-species coalescent (*BEAST). All times are given in thousand years.

|  | Loci Set 1 |  |  | Loci Set 2 |  |  |
| --- | --- | --- | --- | --- | --- | --- |
|  | HPD Lower | Median | HPD Upper | HPD Lower | Median | HPD Upper |
| **Species Split** |  |  |  |  |  |  |
| *Ph. uvarovi*/*Ph. artvinensis* | 63.1 | 117.8 | 191.2 | 59.9 | 116.3 | 188.2 |
| **Population Split** |  |  |  |  |  |  |
| *Ph. uvarovi* | 28.4 | 59.1 | 99.7 | 28.1 | 61.8 | 103.0 |
| *Ph. artvinensis* | 26.3 | 53.7 | 90.0 | 32.4 | 64.5 | 107.3 |

**Fig. 2.**
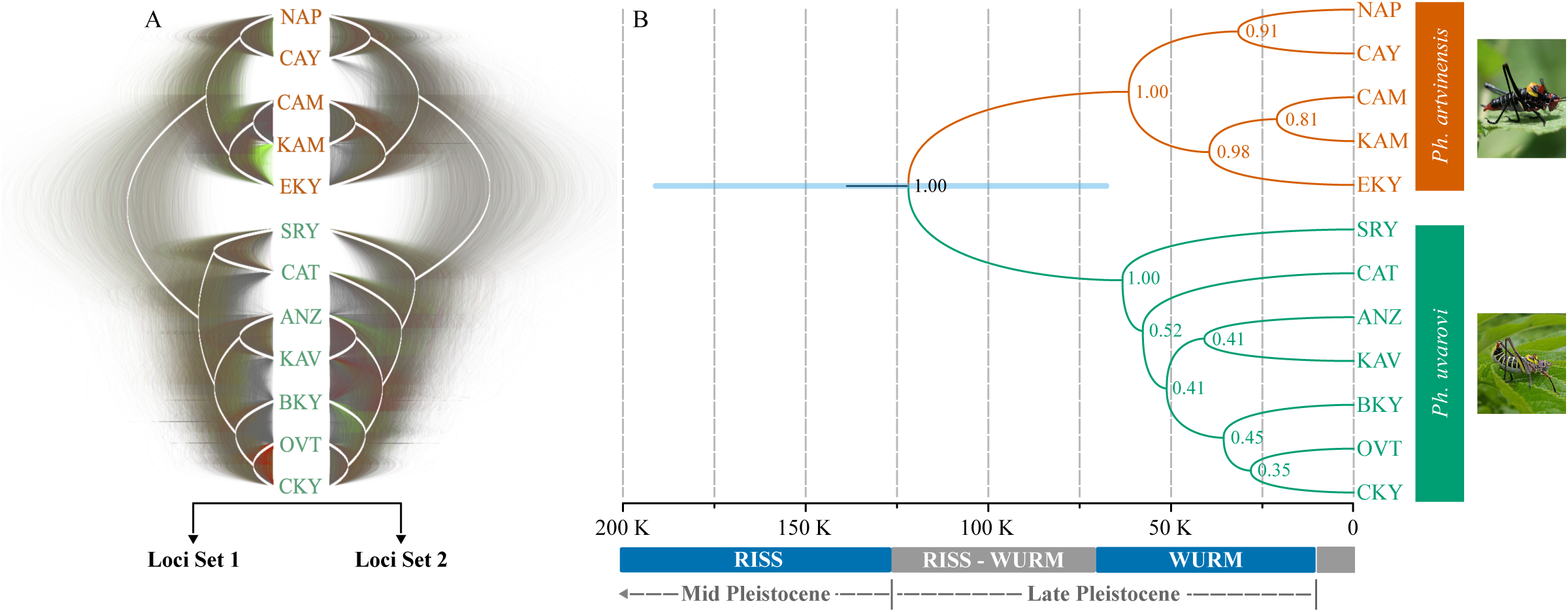
**(A)** Density plots of all trees obtained from *BEAST analyses of the two loci sets. Grey signals describe branching patterns in concert with the consensus topology while green and red indicate increasing degrees of conflict. (**B)** Consensus population tree of *Ph. uvarovi* and *Ph. artvinensis* as summarized over the entire tree space of the two loci sets. Blue bar denotes the 95% highest posterior density interval of the divergence time between the two species.

### 3.3. Different patterns of character displacement for subgenital plate and pronotum of males

In both traits, the “species*type” interaction of the two-way MANCOVA was significant (SGP: Wilk’s λ = 0.808, *F*_*4, 210*_ = 12.435, *P* < 0.001; PRNT: Wilk’s λ = 0.853, *F*_*7, 217*_ = 5.317, *P* < 0.001) but when controlled for population relatedness (pMANCOVA) this interaction was only significant in the subgenital plate (Wilk’s λ = 0.808, *P* < 0.003). For full MANCOVA and pMANCOVA results see Table S4.

The two traits also exhibited differences in the direction of the observed change. The direction of observed change was also different for the two traits. For the subgenital plate the degree of divergence was greater in sympatry vs. allopatry (*D*_*symp-allo*_ = 7.5956; *P*_*rand*_ = < 0.001) whereas for the pronotum divergence in allopatry was greater than in sympatry (*D*_*symp-allo*_ = −3.4922; *P*_*rand*_ < 0.001). We conclude that significant but opposite patterns of character divergence exist in the subgenital plate and pronotum of males, and in the case of the subgenital plate this divergence holds up when controlled for population relatedness.

For patterns in the subgenital plate, both the magnitude (*D*_*mag*_ = 0.225, *P*_*rand*_ = 0.002) and direction (*D*_*dir*_ = 114, *P*_*rand*_ = 0.626) of change exhibited by the two species was clearly different, although results were only significant for the magnitude of change. While *Ph. artvinensis* showed substantial differentiation in shape of the subgenital plate between allopatric and sympatric populations, *Ph. uvarovi* exhibited very little differentiation (Fig. 3A). In contrast, for the pronotum both the magnitude (*D*_*mag*_ = 0.020; *P*_*rand*_ = 0.05) and direction (*D*_*dir*_ = 61.310; *P*_*rand*_ = 0.008) of phenotypic change in allopatry vs. sympatry was similar (Fig. 3B). We conclude that a strong pattern of asymmetric character displacement exists between the two species for divergence of the subgenital plate but not the pronotum.

**Fig. 3.**
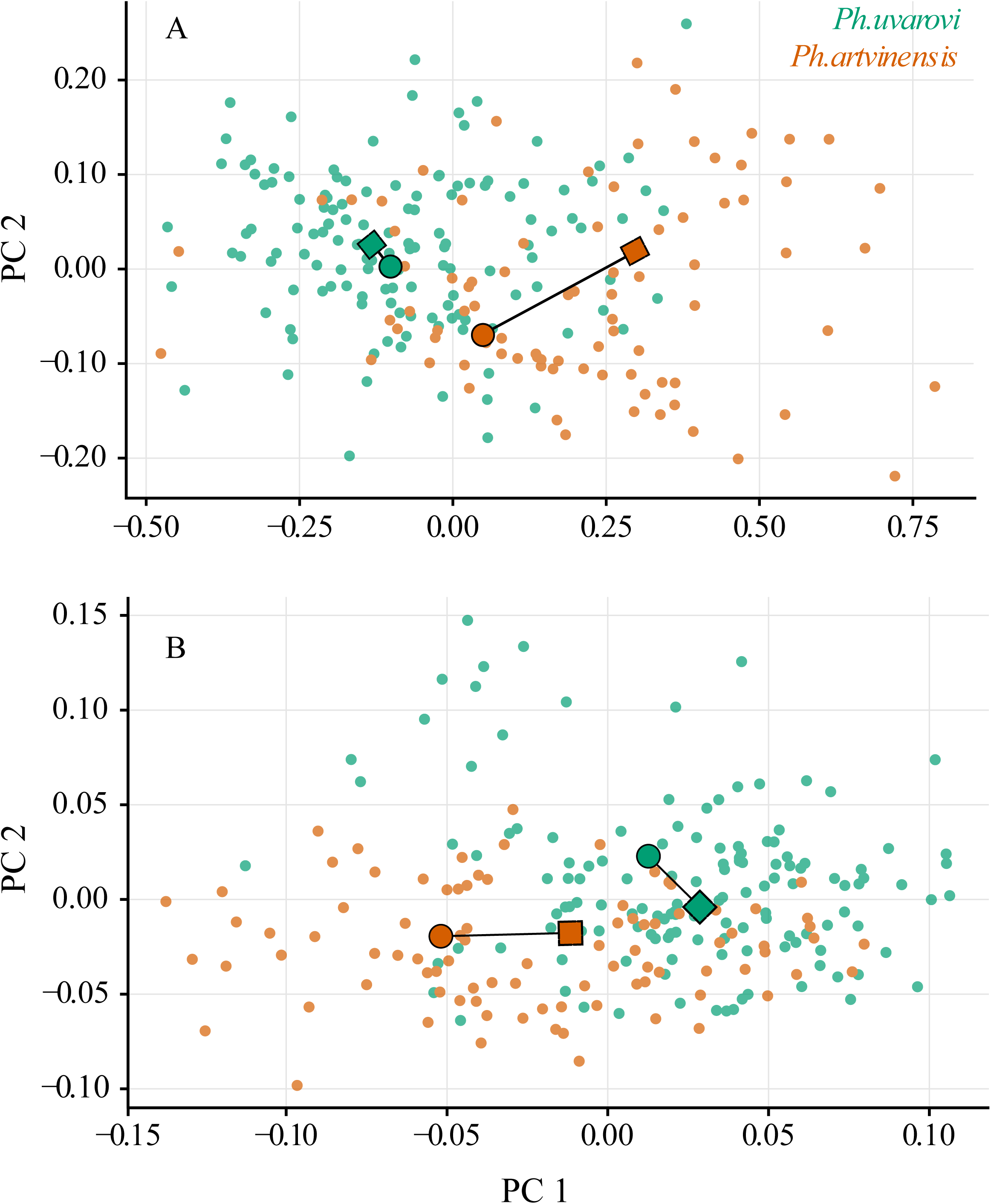
Phenotypic change vectors for the subgenital plate **(A)** and the pronotum **(B)** showing the magnitude and direction of change between *Ph. uvarovi* and *Ph. artvinensis* in allopatry vs. sympatry.

### 3.4. Rapid divergence of the subgenital plate under strong selection and drift

Estimated evolutionary rates indicate the two traits were not evolving at the same rate. In both sets, the likelihood of the unconstrained model (where each trait evolves at a different rate) was substantially higher than the constrained model (Fig. 4A). Moreover, this difference was statistically significant at the 0.001 level for 73% of the tree space in loci set 1 whereas this number rose to 82% for loci set 2 (Fig. S4). More strikingly, our results showed that the evolutionary rate of the subgenital plate was substantially higher than the pronotum with approximately 10-fold differences between median values (Fig. 4B). We conclude that the faster rate of divergence in the subgenital plate indicate the presence of higher evolutionary pressures on this trait.

**Fig. 4.**
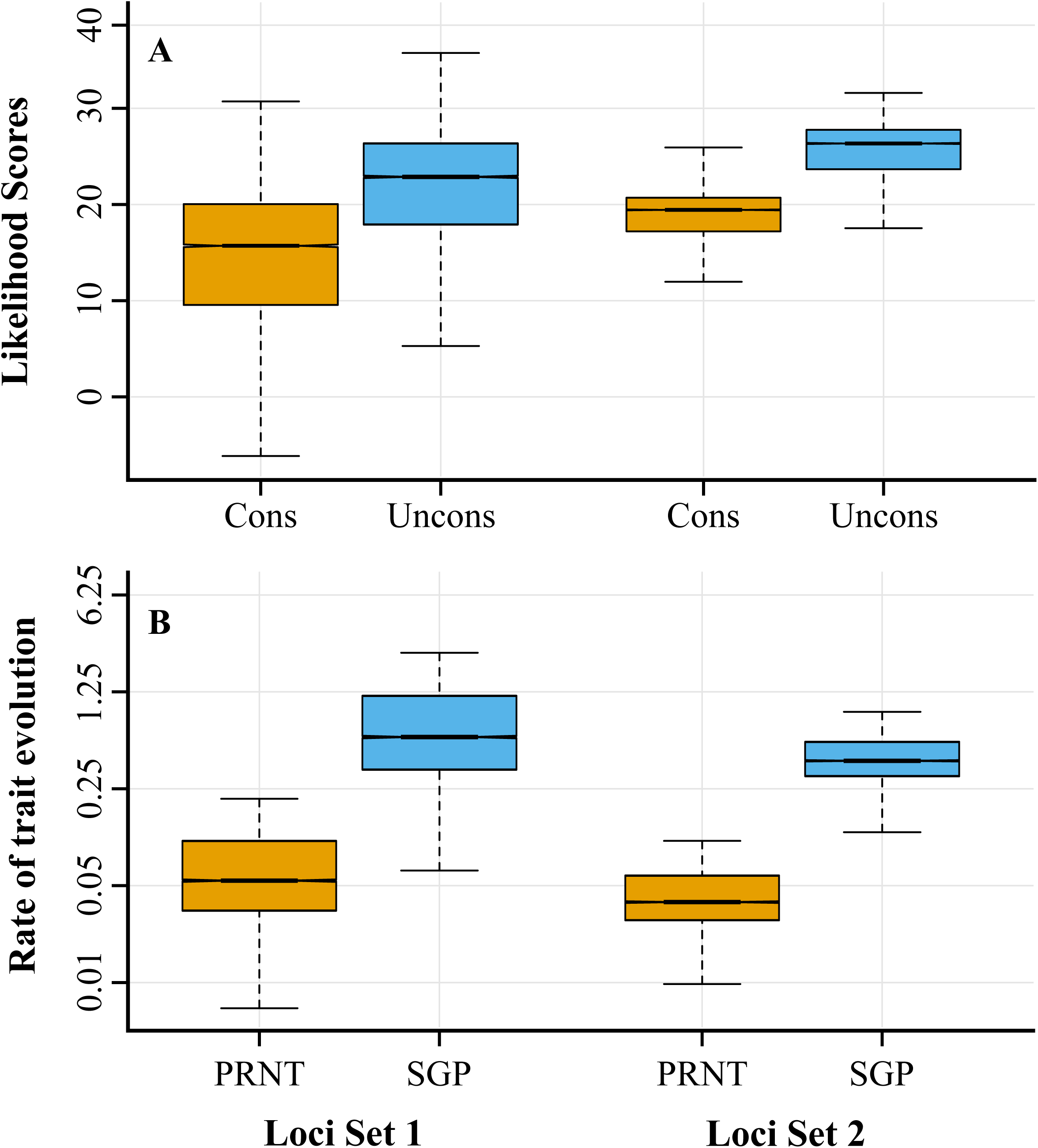
**(A)** Boxplots summarizing the likelihood of constrained and unconstrained models in describing rates of evolution in the two traits (pronotum, subgenital plate). **(B)** Boxplots summarizing estimated evolutionary rates of both traits. Results given separately for the two loci sets.

For divergence in pronotum, likelihood scores did not indicate a significant difference between adaptive (SAP, DAP) and non-adaptive (DM) models (Fig. 5A). Therefore, neutral models of evolution were adequate in explaining difference in divergence of this trait between populations in sympatry vs. allopatry. Strikingly, the double adaptive peak model (DAP) was a substantially better fit than the other two models (DM, SAP) for divergence in the subgenital plate, implying different trait optima for divergence of species in sympatry vs. allopatry (Fig. 5B). The DAP model also returned reasonable optimum trait values for the two selective regimes (Table 2). Moreover, the coefficients of drift and selection for the subgenital plate were substantially higher than optimum trait values (Table 2) indicating strong drift and selection. We conclude that patterns of character displacement in the subgenital plate resulted from adaptation to different trait optima in sympatry and allopatry under both strong drift and selection.

**Table 2.** Parameter estimates under the Double Adaptive Peak (DAP) model of evolution for the subgenital plate. **(σ)** coefficient of drift; **(α)** coefficient of selection; **(*θ***_***allo***_) optimum trait value in allopatry; and **(*θ***_***symp***_) optimum trait value in sympatry.

| Parameter | Loci Set 1 |  |  | Loci Set 2 |  |  |
| --- | --- | --- | --- | --- | --- | --- |
|  | -95% CI | Median | +95% CI | -95% CI | Median | +95% CI |
| $\sigma$ | 55.441 | 57.147 | 59.128 | 73.155 | 74.657 | 75.459 |
| $\alpha$ | 64.587 | 69.100 | 74.069 | 112.399 | 116.773 | 119.221 |
| $\theta_{allo}$ | 15.026 | 15.026 | 15.026 | 15.026 | 15.026 | 15.026 |
| $\theta_{symp}$ | 24.967 | 24.967 | 24.967 | 24.967 | 24.967 | 24.967 |

**Fig. 5.**
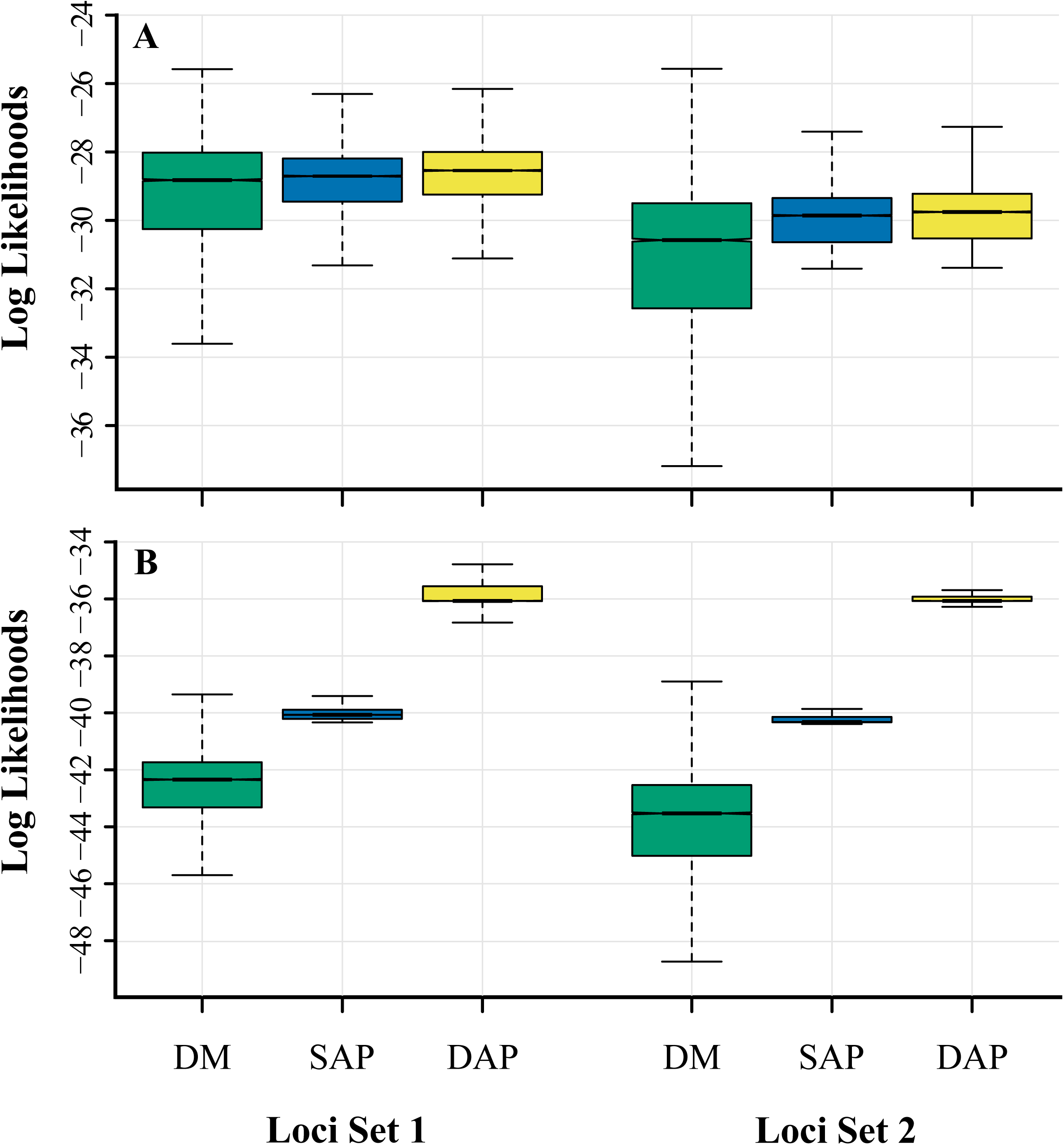
Boxplots summarizing the likelihood of three models: Drift (DM); Single Adaptive Peak (SAP); and Double Adaptive Peak (DAP) in explaining the divergence of the pronotum **(A)** and the subgenital plate **(B)** as calculated separately for the two loci sets.

## 4. Discussion

### 4.1. Character displacement and asymmetric divergence patterns

RCD is one of few processes where natural selection can directly result in reproductive isolation and speciation (Servedio and Noor, 2003; Lukhtanov et al., 2005; Rice and Pfennig, 2010; Yukilevich, 2012). Therefore, reporting patterns of RCD is valuable as it can indicate selection has directly influenced reproductive isolation. When we look at patterns of character divergence between *Ph. uvarovi* and *Ph. artvinensis* we can see that the magnitude of divergence in the reproductive trait was substantially greater in sympatry than allopatry, whereas the opposite was true for the non-reproductive character. This can be taken as an indication that evolution has selectively enhanced reproductive isolation in sympatry in contrast to other forces of local adaptation (Jang et al., 2009).

Asymmetric patterns of character divergence can also be good indicators of selection because species usually differ in competitive ability (Grant, 1972; Schluter, 2000; Hill, 2003) and the cost of reproductive interferences are rarely the same (Lemmon et al., 2004; Pfennig and Stewart, 2011). Both the ECD and RCD literature are full of documented cases where one taxon shows more displacement than the other (Smadja and Ganem, 2005; Cooley et al., 2006; Dingle et al., 2010; Reifová et al., 2011; Bath et al., 2012; Gerhardt, 2013). Such patterns of asymmetric divergence were also observed in our study as higher divergence of species in sympatry was recorded in only one species (i.e. *Ph. artvinensis*). However, because patterns can often be misleading it is important to quantify the role of natural selection in divergence before drawing strong conclusions.

Here we show that phylo-comparative methods in conjunction with genomic data can be useful in filling this gap, even for closely related species/populations.

### 4.2. Importance of accurate evolutionary histories and modelling phylogenetic uncertainty

Although the comparative method can be an extremely powerful tool, one drawback is its heavy reliance on the estimated phylogeny, and on this phylogeny accurately capturing both the topology and evolutionary distances between groups (Boettiger et al., 2012). Phylogenetic inferences based on a few loci usually have low resolution when it comes to separating closely related species or populations. Such low resolution stems from well known factors such as incomplete lineage sorting, polytomies and differences in gene divergence vs. species/population divergence (Knowles and Chan 2008; Sağlam *et al*., 2017). Such concerns have kept phylo-comparative methods from being applied at the population level and have limited their use to deeper, well resolved phylogenies.

Here, we show how modern genomic data in conjunction with coalescent modeling has the power to reconstruct reliable population level phylogenies allowing phylo-comparative methods to be applied and evolutionary parameters estimated. We found that phylogenetic data sets constructed by sampling random genes resulted in well resolved population trees capable of separating populations of species that diverged roughly 120 ka ago (Table 1, Fig. 2). Although the estimated population trees were well resolved, each data set still had a high amount of uncertainty stemming from the variance in gene trees. Therefore, when estimating evolutionary parameters from such population trees it is important to take phylogenetic uncertainty into account. We quantified this uncertainty by using the entire range of sampled trees for conducting phylo-comparative analyses. Furthermore, by conducting all tests under different phylogenetic data sets, we also tested the repeatability of our estimates under different sets of loci. Our approach revealed that parameter estimates were highly influenced by phylogenetic uncertainty as these estimates covered a wide range of values. However, results were fairly consistent under different sets of loci, indicating captured trends were highly repeatable and overall informative.

### 4.3. Rates and adaptive models of trait divergence

In zones of secondary contact, selection to minimize reproductive interactions should exert higher pressures on reproductive versus non-reproductive characters. Therefore in such areas reproductive traits could be expected to show elevated rates of divergence and different adaptive optima in sympatry vs. allopatry. Our results confirm this assumption by showing much faster rates of divergence in the reproductive trait (Fig. 4) and the existence of different selective optima in sympatry vs. allopatry (Fig. 5). On the other hand, the null model of neutral divergence could not be rejected for the non-reproductive trait, presenting further evidence that selection has directly targeted reproductive isolation.

In insects, differences in genital form inflict a high cost on heterospecific matings (Sota and Kubota, 1998; Usami et al., 2006; Sota and Tanabe, 2010) and selection for rapid divergence of genital characters in sympatry relative to allopatry is common (Kawano, 2003; Jang et al., 2009; Kawakami and Tatsuta, 2010). While our results indicate the clear role of selection in rapid character divergence of the reproductive trait, they also highlight the substantial role genetic drift has played in this divergence. Indeed, estimates from both loci sets show the coefficient of drift for the reproductive trait was as high as the coefficient of selection (Table 2). Thus the high amount of displacement observed in the genital trait was only possible under the combination of strong drift and selection. These results can also help determine under which conditions distributional shifts and fragmentation of species during the Quaternary could lead to species level diversification. Such systems are continually characterized by high genetic drift resulting from fragmentation and strong bottlenecks (Knowles and Richards, 2005). We show that when these effects are coupled with strong selection, species level differences could occur even in a relatively short amount of time (i.e. 200 – 70 ka).

## Conclusion

In this paper we provide evidence for many of the criteria needed to infer patterns of diversification between *Ph. uvarovi* and *Ph. artvinensis* resulted from RCD (i.e. from selection acting directly to minimize reproductive interferences). Most notably we document, relatively higher divergence rates, selection towards multiple phenotypic optima and strong asymmetries between species for diversification of the reproductive trait. The patterns and processes outlined in this paper support a model of evolution consistent with allopatric divergence followed by range expansions and secondary contact. A model claimed to have played a prominent role in Anatolian diversity and around the world during climatic shifts of the Quaternary (Hoskin et al., 2005; Çıplak, 2008; Galbreath et al., 2010; Bilgin, 2011). In addition, by quantifying evolutionary parameters from genomic data using phylo-comparative methods we show that both strong drift and selection is necessary to obtain species level diversification via RCD. These results carry important information for bridging the gap between patterns and processes underlying species diversification and for demonstrating the usefulness of the comparative method for quantifying evolutionary hypotheses at the population level.

## 5. Acknowledgements

This research was funded by the Hacettepe University Scientific Research Projects Coordination Unit, Project No’s: 011 D06 601 006-196 and 013 D05 601 005-300. We would very much like to thank Dr. Cem A. Kuyucu, Dr. Çağasan Karacaoğlu and Dr. Duygu P. Öksüz for their valuable help both in the field and laboratory. We would also like to thank Dr. Sibel Küçükyildirim for her help in PCR and molecular protocols, Dr. Bülent Alten and Dr. Özge Erişöz Kasap for laboratory support and Dr. Hasan Sevgili for his valuable input in the systematics and biogeography of the genus.

## Supplementary Material

### Supplementary File 1

#### Methods

Additional details for methods and analyses used within the study.

**Table S1**. Population and sampling information.

**Table S2**. GenBank accession numbers of ITS and mtDNA sequences.

**Table S3**. Principal component decomposition of shape variables from fourier analysis.

**Table S4**. Full MANCOVA results.

**Fig. S1**. Representative images of traits.

**Fig. S2**. Visualization of shape variation.

**Fig. S3**. Phylogenetic representation of alternative models of trait evolution.

**Fig. S4**. Likelihood ratio tests comparing constrained and unconstrained rates of trait evolution.

**Fig. S5**. mtDNA gene tree.

**Fig. S6**. ITS gene tree.

**Fig. S7**. Consensus population tree using only RAD loci from loci set 1.

**Fig. S8**. Consensus population tree using only RAD loci from loci set 2.

### Supplementary File 2

#### Data archive

**F**ile containing all necessary data to reproduce the current study. Contains, full shape data as a csv file, the *de-novo* set of 45,039 reference RAD-contigs, phylogenetic data sets in FASTA format, XML files used in BEAST analyses and individual gene trees in nexus format.

