## Supplementary material for "Phylo-comparative analyses reveal the dual role of drift and selection in reproductive character displacement"

### Additional methods

#### *DNA extraction and sequence data generation*

DNA was extracted from the hind femura of 6 samples from each of the 12 populations using E.Z.N.A.® Insect DNA Kit (Omega Bio-Tek) according to the manufacturer's protocol. High throughput genomic data was generated by paired-end RAD sequencing and RAD libraries were constructed using the Sbf1 eight base recognition site restriction enzyme and new RAD protocol described in (Ali *et al.*, 2016). In total we generated 347,347,272 sequence pair reads across all 72 individuals with an average sequence depth of 4X.

ITS1 & ITS2 regions were amplified from 3 individuals from each of the 12 populations using the primer set 18S and 28S (Weekers *et al.*, 2001) and following the PCR protocol given in (Ullrich *et al.*, 2010). Sequencing reactions were performed by Macrogen (908 World Meridian Centre #60-24; Gasan-dong, Geumchon-gu Seoul, Korea, 153-023) on an Applied Biosystems 3730xl DNA sequencer using the amplifying primers. Sequences were proof read using both forward and reverse primers and were aligned in Clustal X2 (Larkin *et al.*, 2007). The front and end portions of sequences were trimmed to ensure that all sequences were of the same length and final sequence length of the combined ITS 1 and ITS 2 regions was 773 bp.

mtDNA sequences were obtained from the concatenated data set given in (Sağlam *et al.*, 2014) comprised of 582 bp segment of the ND4 gene, a 312 bp segment of the Cytb gene and a 108 bp segment of the ND1 (total length 1002 bp).

#### *De-novo RAD-loci discovery and partial genome reference*

The generated sequences were sorted into individuals using 8 bp unique barcodes. Reads containing exact matches to the 8 bp barcodes followed by the 8 bp cut site were sorted into separate files and the 16 bp identifier sequence was removed. For each file, matching reads pairs were determined using header information and written to separate files. *De-nov*o loci discovery and extension using read pairs was carried out using custom PERL scripts, the alignment program Novoalign and the genome assembler PRICE (Ruby *et al.*, 2013).

RAD-loci discovery was conducted using six individuals from the CAT population of *Ph. uvarovi* with the procedure outlined in (Miller *et al.*, 2012). In short, the 88 bp forward reads were trimmed from the 3' end to 80 bp, quality filtered using a cut-off of 20% and concatenated into a single FASTA formatted file. This file was then indexed using Novoalign and aligned to its own index. This produced an alignment map containing pairwise alignment scores of internal (within individuals) and external (between individuals) alignments. Using this map we determined distinct loci based on the following criteria: The maximum alignment score between any sequences in a locus was 30, alignments with a score over 90 and perfect internal alignments were ignored; sequences that had only one occurrence were ignored unless they had a perfect external alignment with a sequence that had more than one occurrence; each described locus had to contain one sequence from at least two individuals.

Using the above procedure, we were able to identify 75,473 putative RAD-loci. These RAD-loci were then extended into longer contigs using the genome assembler PRICE (Ruby *et al.*, 2013) which used paired end read information to iteratively increase existing contigs. Each putative RAD locus was extended in PRICE separately. For each locus we created 3 files: 1) A template file containing the 80 bp RAD-locus with the cut site added back on; 2) A forward read file containing all exact matches to the 80 bp RAD-locus and 3) A reverse read file containing the corresponding paired reads. In addition, we added a 100 bp poly A block in front of the cut site to ensure PRICE would not extend upstream of the cut site. These three files were then entered into PRICE and the existing RAD-locus extended using the following settings: the insert size and percent identity required for aligning a read to a contig was set at 300 and 95% respectively; the minimum overlap length for extension was set to 40 nt (-mol 40); the minimum percent identity for a sequence to match was set at 80 nt (-mpi 80). PRICE was run on 100% target mode (-target 100 0) and the cycle number was kept at 1 (-nc 1), forcing PRICE to only extend the input contigs.

Using the above procedure, we were able to extend 45,039 of the 75,473 putative RAD-loci into contigs ranging between 300-800 bp. RAD-loci which failed to extend and contigs shorter than 300 bp were discarded. The final set of 45,039 RAD-contigs were used as a *de-novo* partial genome reference for all subsequent downstream analyses.

***Phylogentic data sets and analysis***

Individuals were aligned against the reference set of RAD-contigs (partial reference genome) using the BWA-MEM algorithm (Li & Durbin, 2009; Li, 2013) and outputted to BAM file format. BAM files were then filtered to remove read pairs that did not align to the same sequence and those representing PCR duplicates using SAMtools (Li *et al.*, 2009). Resulting BAM files were then indexed and used in obtaining consensus sequences for each individual. Consensus sequences were built by using individual read depths to call for the most common base at a site using the probabilistic framework in ANGSD (Datta & Nettleton, 2014). Bases with a quality score below 13 on the phred 33 scale were disregarded. In case of ties a random base was chosen from among the bases with the highest counts. Ambiguous sites due to overall low read quality and depth was outputted as "N". From our set of reference loci we then randomly chose 1000 loci for phylogenetic analysis and filtered out any loci that did not contain a sample from at least three individuals of a given population. From the remaining set of loci we conducted two random draws of 50 RAD-loci without replacement to build two independent genomic subsets. To each subset we further added all available mtDNA and ITS sequences of each population to come up with two phylogenetic data sets, each comprised of 52 loci.

Prior to phylogenetic analysis we determined the best-fit model of evolution for each loci independently using the Akaike and Bayesian Information criterions as implemented in MEGA6 (Tamura *et al.*, 2013). All RAD loci best confirmed with the JC (Jukes-Cantor) model of evolution (Jukes & Cantor, 1969) under both criteria. For the mtDNA and ITS data sets the most appropriate models of evolution were found to be the HKY+G (Hasegawa *et al.*, 1985) and JC + G model respectively. For each loci set we built separate Xml files in BEAUTI version 2.2.0 for analysis in

BEAST. All analysis were set up using a strict molecular clock, a linear population size model for the multi-species coalescent and a Yule model of divergence. Trees were time calibrated by applying a log-normal prior on the mtDNA clock rate parameter with a mean of 0.0133 ± 0.0013 (subs/s/my/l) in real space. This calibration corresponds to a global mtDNA rate of 2.6% divergence per Myr which strikes a balance between the faster rates of cytb and slower rates of Nad4 and Nad1 used within this study (for further information please refer to) (Sağlam *et al.*, 2014). Clock rate parameters of all remaining loci (i.e. RAD loci + ITS) were drawn from a uniform distribution between 0 – 100. For the birth rate parameter (λ) of the Yule model we used a uniform prior between 0 – 1000 whereas for the population size model we used the Jeffrey's prior which is a scale invariant uninformative prior [15].

***Independence of Phylogenetic sets and resulting population trees***

Since both phylogenetic sets contained mtDNA and ITS regions, questions might arise whether robustness of resulting population trees are driven by the inclusion of common loci or whether the two sets are truly independent. To clarify these issues, we present mtDNA and ITS gene trees below along with population trees reconstructed using only RAD loci.

Phylogenetic inferences from mtDNA trees were unable to resolve evolutionary history of populations and showed extensive haplotype sharing between *Ph. artvinensis* and *Ph. uvarovi* (Fig. S5). Trees built using ITS regions were able to resolve species but could not distinguish between populations (Fig. S6). Thereby it is not likely that similarity between estimated population trees using the full data set are driven by shared loci. Moreover, phylogenetic patterns obtained from the two RAD-loci sets after removal of shared loci (Fig. S7-8) were consistent with trees obtained from the full data sets. This shows that phylogenetic inferences were robust and the two sets were independent despite sharing mtDNA and ITS markers. Thereby, both mtDNA and ITS regions were kept in for calibrating the tree as mutation rates for these loci are relatively well known.

**Supplementary tables**

**Table S1. Population information**. Species affiliation (artv = *Ph. artvinensis*; uvar = *Ph. uvarovi*), abbreviations of population names (**Codes**), altitude (**Alt**), **Type** (allopatric/sympatric), number of individuals used in shape measurements (**N**), mean size of each trait (trait area as measured in number of pixels), mean PC1 scores and measurement errors (**ME**) of shape variation and the average mahalanobis distance (**Mahb**) of each population from like populations of the other species (i.e. allopatric vs allopatric and sympatric vs sympatric) are given.

| Population | Species | Codes | Alt (m) | Type | Subgenital Plate | | | | | Pronotum | | | | |
|---|---|---|---|---|---|---|---|---|---|---|---|---|---|---|
| | | | | | N | Size | PC1 | ME | Mahb | N | Size | PC1 | ME | Mahb |
| Erikli Köyü (1) | artv | EKY | 1802 | Symp | 18 | 405259 | 0.348 | 0.041 | 22.829 | 18 | 299135 | -0.019 | 0.001 | 14.678 |
| Boğalı Köy (2) | uvar | BKY | 2139 | Symp | 18 | 311045 | -0.127 | 0.025 | 33.423 | 20 | 246493 | -0.031 | 0.001 | 7.519 |
| Anzer Yaylası (3) | uvar | ANZ | 2154 | Symp | 19 | 305970 | -0.128 | 0.022 | 22.299 | 20 | 231340 | -0.045 | 0.001 | 4.064 |
| Kama Köyü (4) | artv | KAM | 1432 | Symp | 11 | 396225 | 0.360 | 0.038 | 29.636 | 14 | 271327 | 0.040 | 0.002 | 12.905 |
| Ovit Geçidi (5) | uvar | OVT | 1857 | Symp | 20 | 301126 | -0.035 | 0.027 | 17.218 | 20 | 247825 | -0.028 | 0.001 | 6.203 |
| Çamlık Köyü (6) | artv | CAM | 1484 | Symp | 20 | 381481 | 0.285 | 0.036 | 24.399 | 17 | 280062 | 0.002 | 0.002 | 8.748 |
| Çağrankaya Yaylası (7) | uvar | CKY | 2226 | Allo | 19 | 283738 | -0.074 | 0.041 | 17.731 | 17 | 278296 | 0.038 | 0.001 | 5.707 |
| Alt Çat (8) | uvar | CAT | 1246 | Allo | 20 | 349984 | 0.113 | 0.033 | 16.013 | 20 | 254206 | -0.016 | 0.001 | 10.312 |
| Kavron Yaylası (9) | uvar | KAV | 1982 | Allo | 20 | 281563 | -0.141 | 0.012 | 8.297 | 20 | 228857 | -0.051 | 0.003 | 6.059 |
| Çaymakçur Yaylası (10) | artv | CAY | 1996 | Allo | 20 | 375847 | -0.024 | 0.040 | 9.491 | 20 | 261022 | 0.015 | 0.002 | 11.699 |
| Sırt Yaylası (11) | uvar | SRY | 2346 | Allo | 19 | 262805 | -0.174 | 0.032 | 18.791 | 20 | 228037 | -0.027 | 0.001 | 5.911 |
| Napopen Yaylası (12) | artv | NAP | 2171 | Allo | 20 | 383502 | 0.187 | 0.053 | 19.830 | 18 | 307178 | 0.073 | 0.001 | 10.947 |

**Table S2.** GenBank accession numbers of ITS and mtDNA sequences used within the study

| Species | Population | ITS1 | ITS2 | mtDNA ND4 | mtDNA cytb-ND1 |
|---|---|---|---|---|---|
| | | KT967141 | KT967177 | KF686939 | KF686885 |
| | CAT01 | KT967142 | KT967178 | KF686940 | KF686886 |
| | | KT967143 | KT967179 | KF686941 | KF686887 |
| | | KT967144 | KT967180 | KF686948 | KF686894 |
| | ANZ01 | KT967145 | KT967181 | KF686949 | KF686895 |
| | | KT967146 | KT967182 | KF686950 | KF686896 |
| | | KT967147 | KT967183 | KF686963 | KF686906 |
| | CKY01 | KT967148 | KT967184 | KF686964 | KF686907 |
| | | KT967149 | KT967185 | KF686965 | KF686908 |
| | | KT967150 | KT967186 | KF686984 | KF686924 |
| ***Ph.uvarovi*** | KAV01 | KT967151 | KT967187 | KF686985 | KF686925 |
| | | KT967152 | KT967188 | KF686986 | KF686926 |
| | | KT967153 | KT967189 | KF686993 | KF686930 |
| | OVT01 | KT967154 | KT967190 | KF686994 | KF686931 |
| | | KT967155 | KT967191 | KF686995 | KF686932 |
| | | KT967156 | KT967192 | KF686951 | KF686897 |
| | BKY01 | KT967157 | KT967193 | KF686952 | KF686898 |
| | | KT967158 | KT967194 | KF686953 | KF686899 |
| | | KT967159 | KT967195 | KF686996 | KF686933 |
| | SRY01 | KT967160 | KT967196 | KF686997 | KF686934 |
| | | KT967161 | KT967197 | KF686998 | KF686935 |
| | | KT967162 | KT967198 | KF686957 | KF686900 |
| | CAM01 | KT967163 | KT967199 | KF686958 | KF686901 |
| | | KT967164 | KT967200 | KF686959 | KF686902 |
| | | KT967165 | KT967201 | KF686960 | KF686903 |
| | CAY01 | KT967166 | KT967202 | KF686961 | KF686904 |
| | | KT967167 | KT967203 | KF686962 | KF686905 |
| | | KT967168 | KT967204 | KF686975 | KF686918 |
| ***Ph. artvinensis*** | EKY01 | KT967169 | KT967205 | KF686976 | KF686919 |
| | | KT967170 | KT967206 | KF686977 | KF686920 |
| | | KT967171 | KT967207 | KF686942 | KF686888 |
| | KAM01 | KT967172 | KT967208 | KF686943 | KF686889 |
| | | KT967173 | KT967209 | KF686944 | KF686890 |
| | | KT967174 | KT967210 | KF686987 | KF686927 |
| | NAP01 | KT967175 | KT967211 | KF686988 | KF686928 |
| | | KT967176 | KT967212 | KF686989 | KF686929 |

**Table S3.** Principle component decomposition of shape variables obtained from fourier analysis. Eigenvalues and percent of variation explained by each axes are shown. Only PC axes summarizing a significant portion of the variation are given.

| Component | Subgenital Plate | | | Pronotum | | |
|---|---|---|---|---|---|---|
| | **Eigenvalue** | **Proportion (%)** | **Total (%)** | **Eigenvalue** | **Proportion (%)** | **Total (%)** |
| **PC1** | 0.0675 | 79.8021 | 79.8021 | 0.0029 | 45.8560 | 45.8560 |
| **PC2** | 0.0081 | 9.6249 | 89.4270 | 0.0016 | 25.3395 | 71.1955 |
| **PC3** | 0.0042 | 4.9647 | 94.3917 | 0.0006 | 9.9242 | 81.1197 |
| **PC4** | 0.0015 | 1.8014 | 96.1931 | 0.0005 | 7.2314 | 88.3510 |
| **PC5** | | | | 0.0002 | 2.8212 | 91.1722 |
| **PC6** | | | | 0.0001 | 1.6953 | 92.8676 |
| **PC7** | | | | 0.0001 | 1.6152 | 94.4828 |
| **PC8** | | | | 0.0001 | 0.9284 | 95.4112 |
| **PC9** | | | | 0.0000 | 0.6926 | 96.1038 |

**Table S4.** Full table of MANCOVA and pMANCOVA results. Significant PC axes were used as dependant variables. Species and locality (allopatric/sympatric) were entered as fixed factors and size and altitude were entered as covariates.

| Trait | MANCOVA | | | | | *pMANCOVA* | |
|---|---|---|---|---|---|---|---|
| | **Wilk's Λ** | **F** | **Effect** | **Error** | ***P*** | ***Wilk's Λ*** | ***P*** |
| **Pronotum** | | | | | | | |
| Intercept | 0.822 | 6.712 | 7 | 217 | < **0.001** | 0.568 | 0.354 |
| Altitude | 0.969 | 1.001 | 7 | 217 | 0.432 | 0.989 | 0.994 |
| Size | 0.795 | 7.996 | 7 | 217 | < **0.001** | 0.525 | 0.248 |
| Species | 0.610 | 19.847 | 7 | 217 | < **0.001** | 0.275 | 0.067 |
| Type | 0.595 | 21.113 | 7 | 217 | < **0.001** | 0.478 | 0.292 |
| Species*Locality | 0.854 | 5.317 | 7 | 217 | < **0.001** | 0.611 | 0.456 |
| **Subgenital Plate** | | | | | | | |
| Intercept | 0.837 | 10.238 | 4 | 210 | < **0.001** | 0.225 | **0.032** |
| Altitude | 0.833 | 10.559 | 4 | 210 | < **0.001** | 0.379 | 0.134 |
| Size | 0.877 | 7.352 | 4 | 210 | < **0.001** | 0.204 | **0.033** |
| Species | 0.585 | 37.315 | 4 | 210 | < **0.001** | 0.215 | **0.040** |
| Type | 0.752 | 17.347 | 4 | 210 | < **0.001** | 0.119 | **0.010** |
| Species*Locality | 0.808 | 12.436 | 4 | 210 | < **0.001** | 0.156 | **0.003** |

### 3. Supplemantary figures

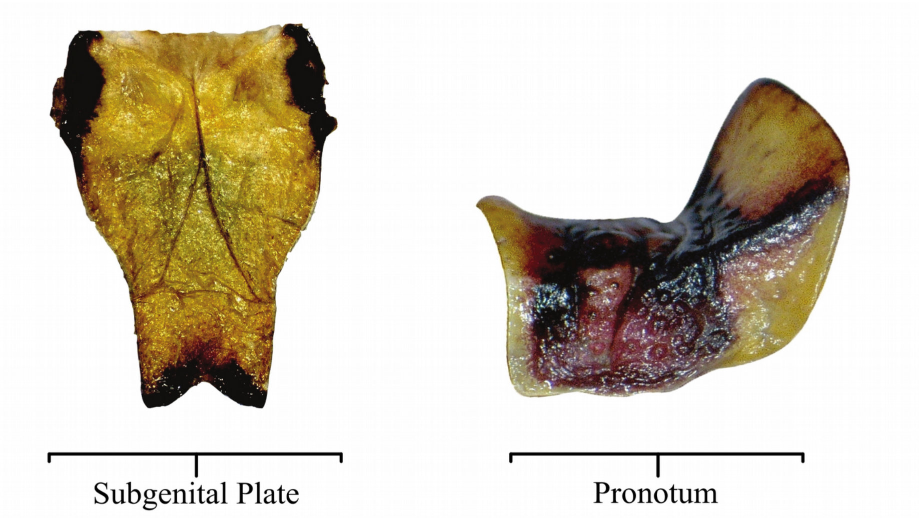


**Figure S1.** Representative images of traits used in phenotypic analysis

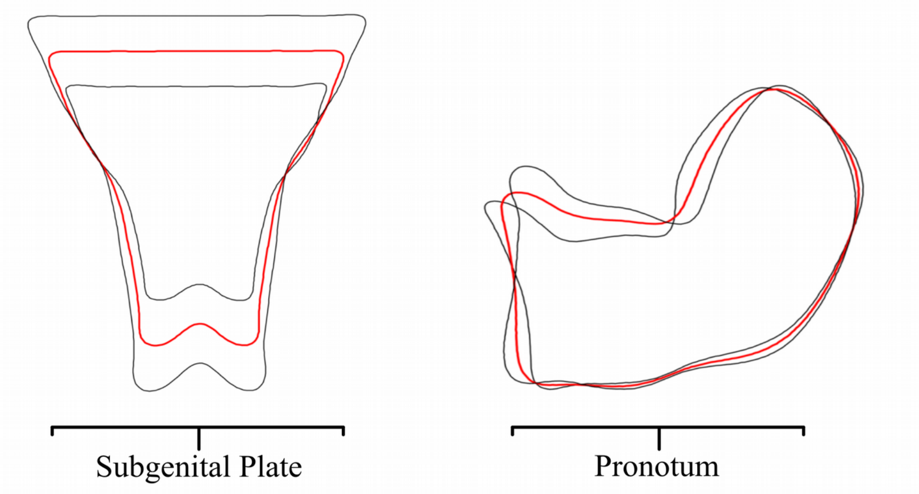


**Figure S2.** Visualization of shape variation described by PC1 for both traits. Red traces represent median shape while black traces represent the upper and lower 95% CI of shape variation.

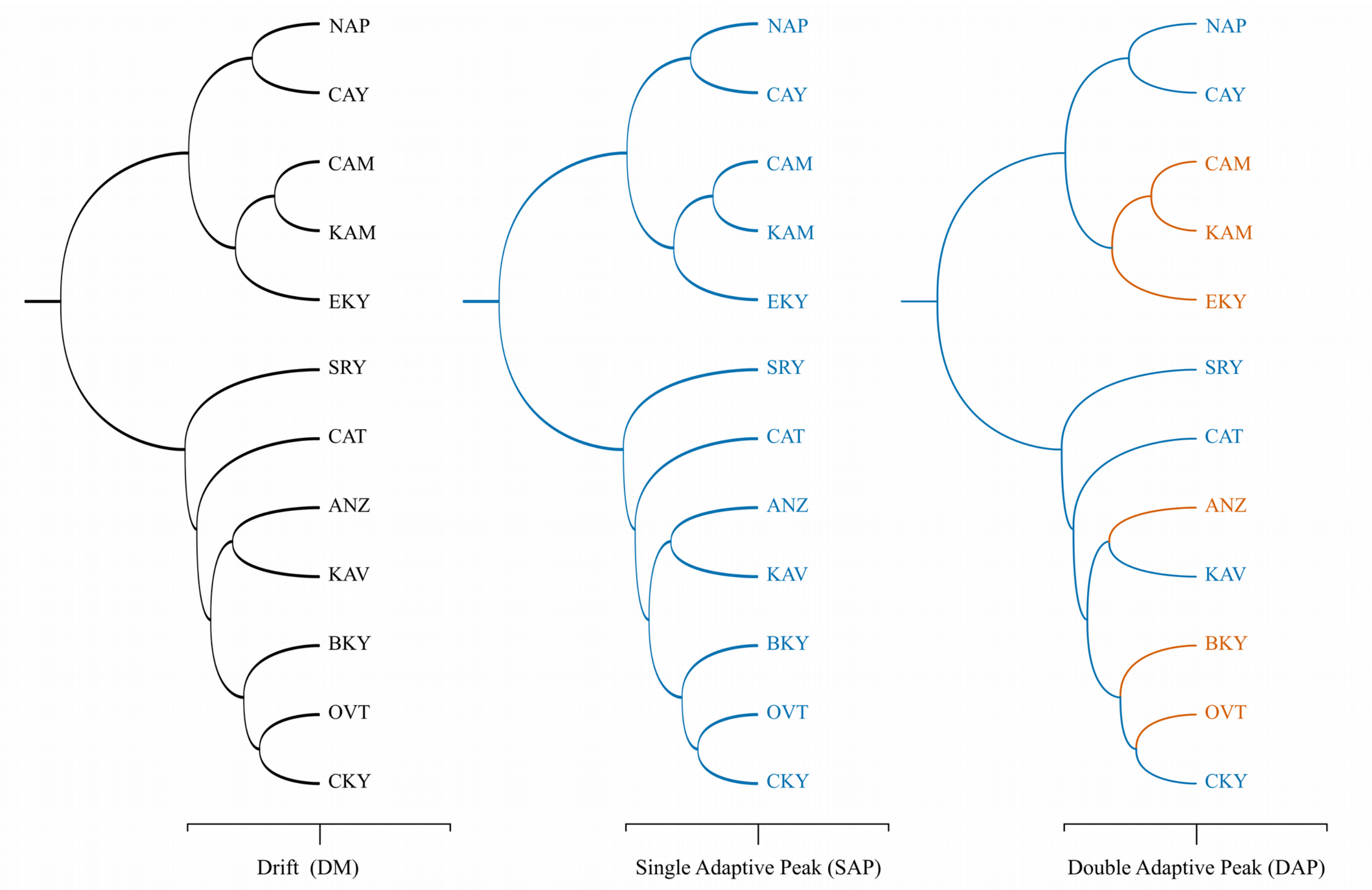


**Figure S3.** Alternative models of trait evolution as visualized on the population tree of *Ph. uvarovi* and *Ph. artvinensi*s. Color codes indicate different adaptive optima for the degree of divergence between species in sympatry vs allopatry. Black for no adaptive optima in the drift only model, blue for a single uniform adaptive peak for all populations in the SAP model and blue and red for different trait optima for populations in sympatry (red) and allopatry (blue) in the DAP model.

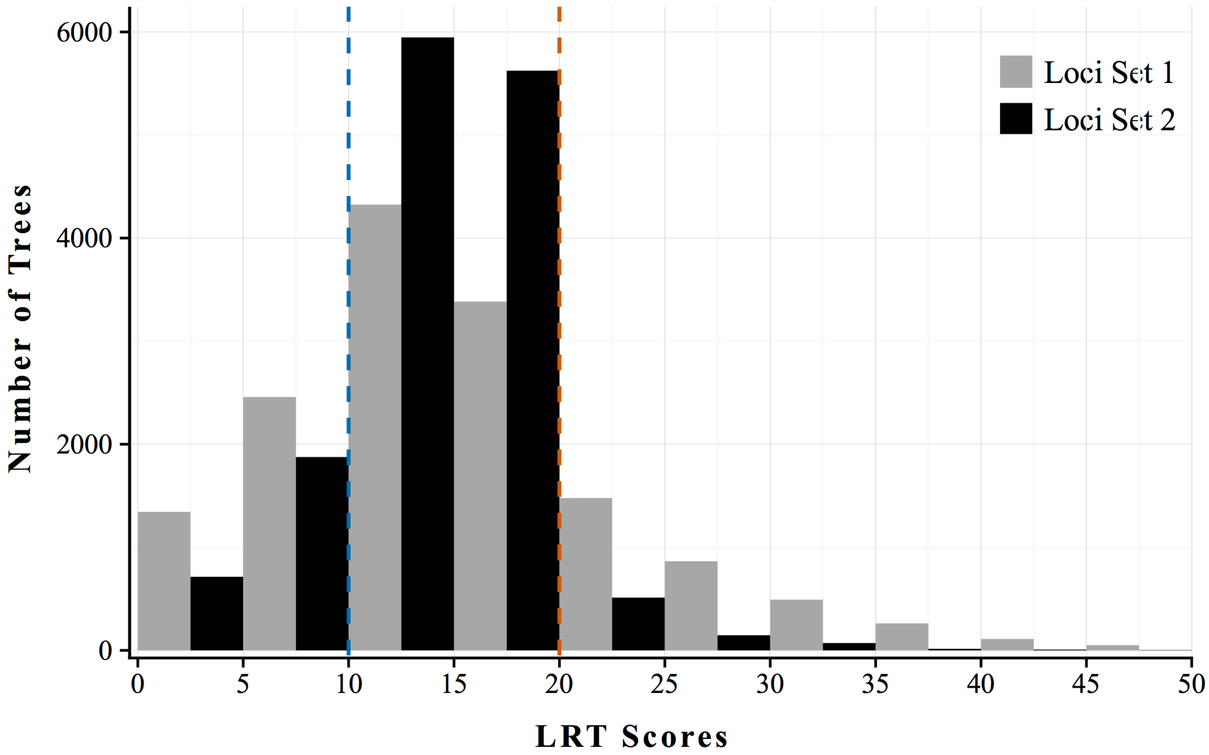


**Figure S4.** Likelihood ratio tests between constrained and unconstrained rates of trait evolution of the pronotum and subgenital plate, as conducted over all trees in the two genomic subsets. Blue dashed line denotes statistical significance at the 0.05 level and red dash line at the 0.001 level.

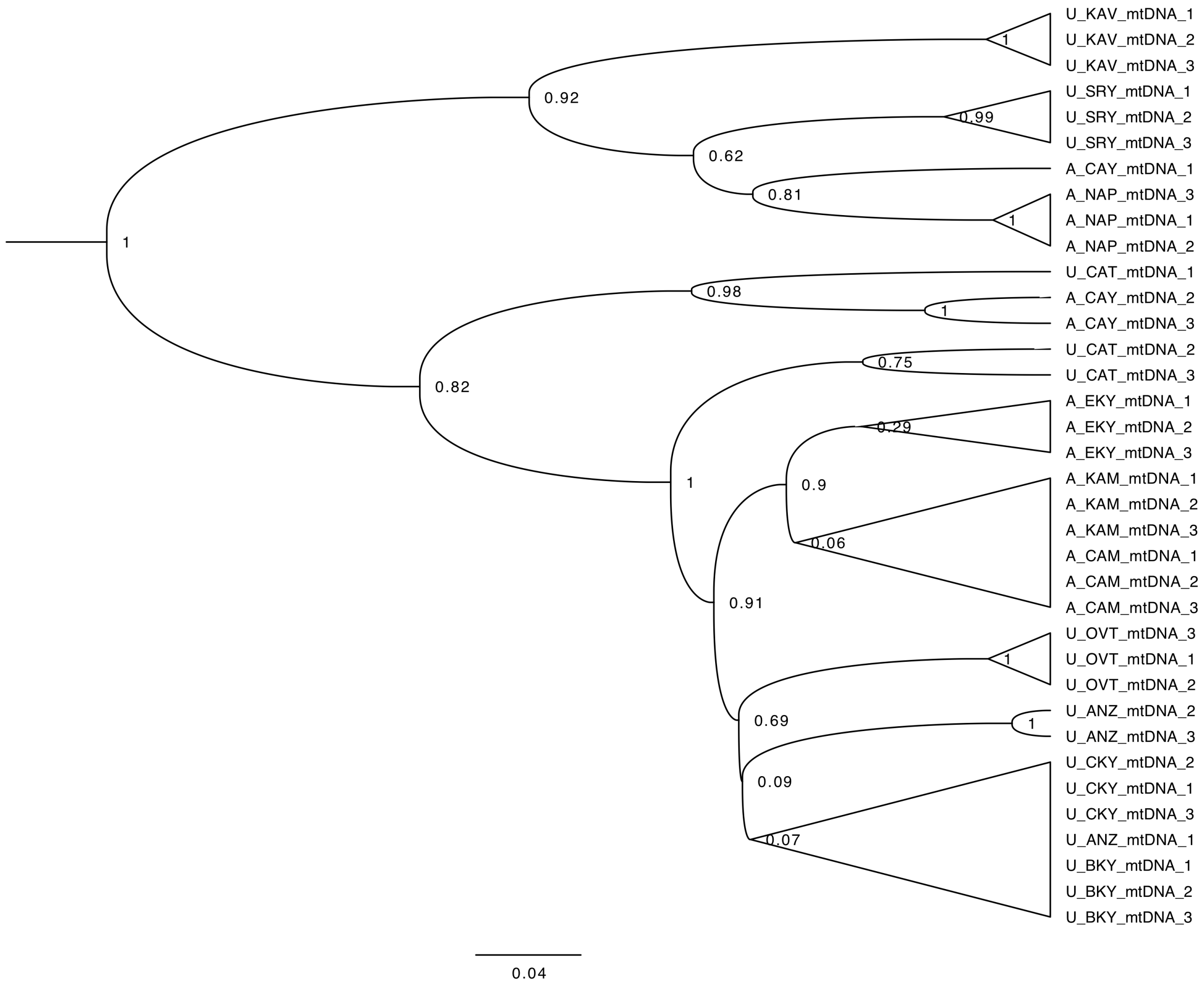


**Fig. S5.** mtDNA gene tree reconstructed from a concatenated data set of partial sequences of ND4 (582 bp), Cytb (312 bp) and ND1 (108 bp) genes. Total length 73 bp.

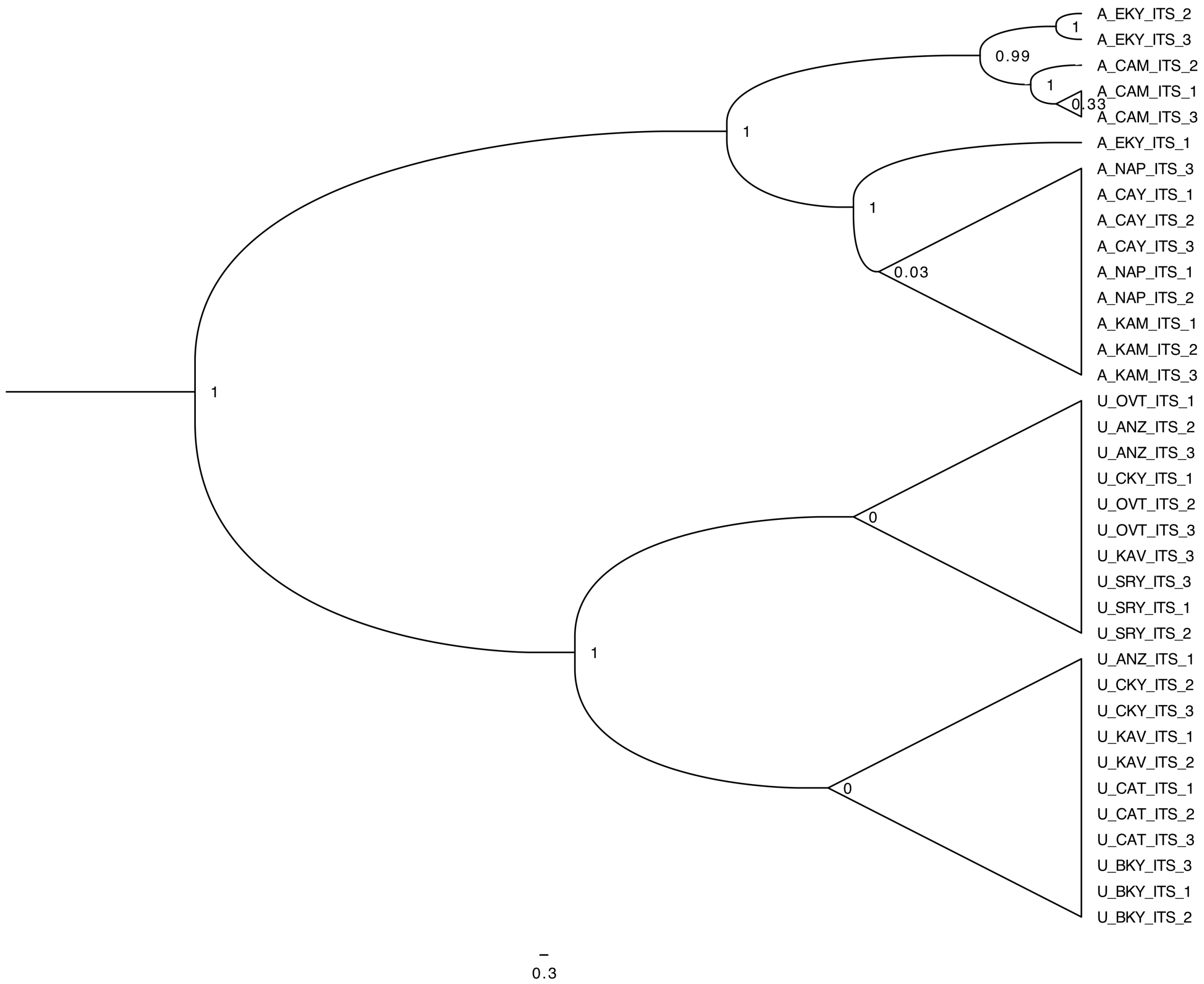


**Fig. S6.** ITS gene tree reconstructed from combined sequences of the full ITS 1 and 2 regions. Total length 773 bp.

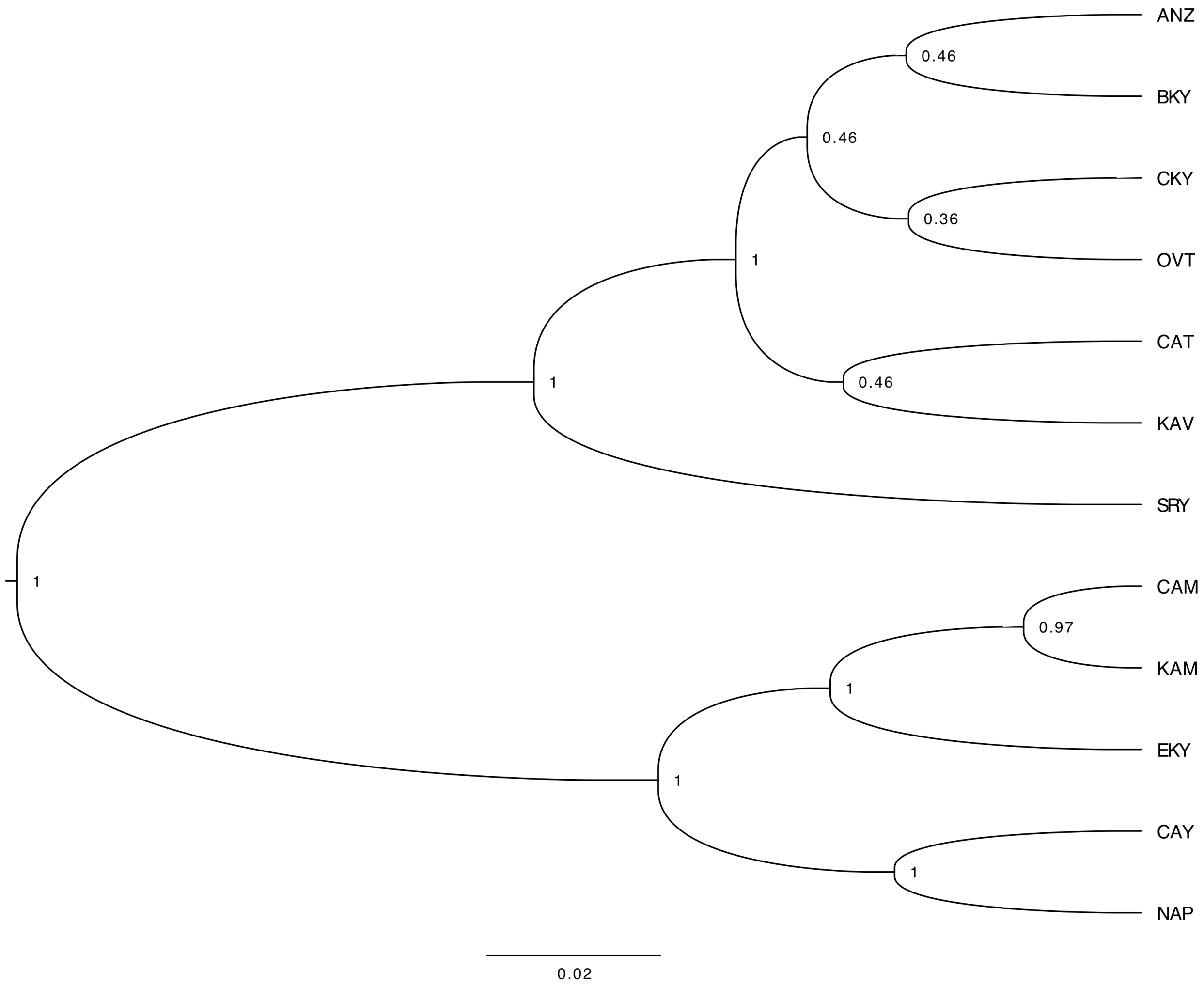


**Fig. S7.** Consensus population tree of Ph. Uvarovi and Ph. Artvinensis as constructed from 50 RAD loci in locus set 1.

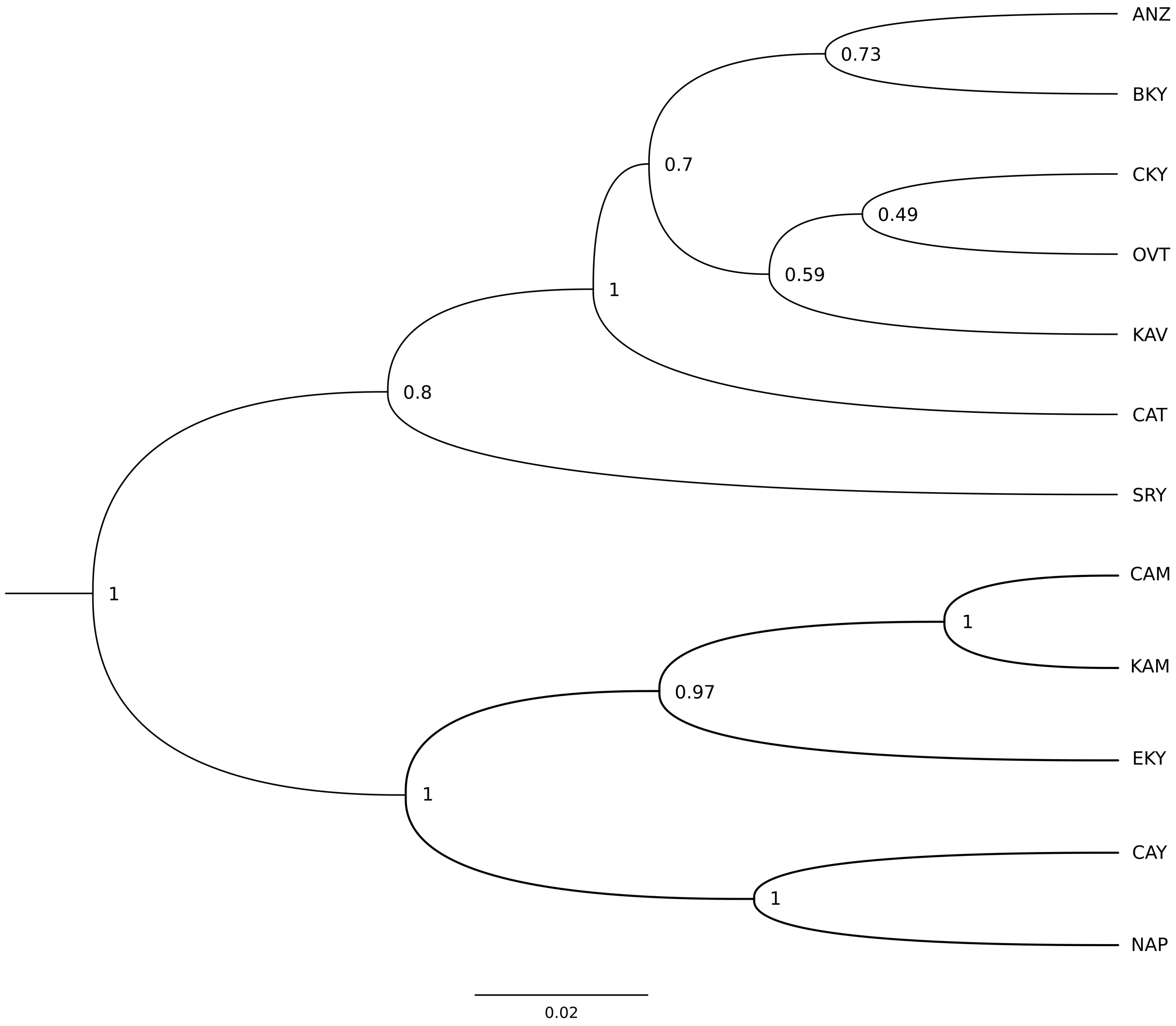


**Fig. S8.** Consensus population tree of *Ph. uvarovi* and *Ph. artvinensis* as constructed from 50 RAD loci in locus set 2.
